# Biocontrol efficiency and mechanism of novel *Streptomyces luomodiensis* SCA4-21 against banana *Fusarium* wilt

**DOI:** 10.1101/2024.10.22.619680

**Authors:** Qiao Liu, Liangping Zou, Yufeng Chen, Junting Feng, Yongzan Wei, Zhang Miaoyi, Kai Li, Yankun Zhao, Dengbo Zhou, Wei Wang, Dengfeng Qi, Jianghui Xie

## Abstract

The soil-borne fungi *Fusarium oxysporum* f. sp. *cubense* tropical race 4 (Foc TR4) causes banana *Fusarium* wilt, which seriously threats global banana production. Biocontrol has been considered viable alternative method to manage banana *Fusarium* wilt. In previous study, we found that novel *S. luomodiensis* SCA4-21 exhibited antifungal activity. Here, we revealed that the extracts of strain SCA4-21 significantly inhibited the growth of Foc TR4 hyphae and spore germination, severely disrupting the ultrastructure of Foc TR4 hyphal cells and spore morphology. These extracts also exhibited broad-spectrum antifungal activity against eight other phytopathogenic fungi. Furthermore, we demonstrated that strain SCA4-21 produce 32 volatile organic compounds, including five antifungal compounds. In a pot experiment, we discovered that the inoculation of strain SCA4-21 significantly inhibited the infection of Foc TR4 in banana seedling corms, achieving a biocontrol efficiency of 59.3%, and promoted the growth of banana seedlings. Additionally, this inoculation significantly enhanced the abundances of beneficial bacterial genera *Streptomyces*, *Bacillus*, *Sphingomonas*, and *Massilia*, as well as fungal genera *Mortierella*, *Purpureocillium*, *Gibellulopsis*, and *Xenomyrothecium*, while significantly reduced the abundances of pathogenic bacteria genus *Pantoea* and fungal genus *Fusarium*, in the banana rhizosphere soil. Moreover, the inoculation of strain SCA4-21 significantly increased the enrichment of pathways such as carbohydrate metabolism, amino acid metabolism, and metabolism of terpenoids and polyketides. Therefore, we postulated that strain SCA4-21 may synergistically combat banana *Fusarium* wilt by producing antifungal compounds and enriching beneficial bacteria and fungi. Our findings present a promising biocontrol agent for the management of banana *Fusarium* wilt.

**IMPORTANCE:** Banana (*Musa* spp.) is one of the most popular fruit crops and the fourth largest food crop in developing countries within tropical and subtropical regions. However, the emergence and rapid spread of strain Foc TR4 seriously hinder the development of banana industry. Currently, there is no effective control measure available. Biological control holds potential due to its safety and effectiveness. Here, we found that the extracts of *S. luomodiensis* SCA4-21 exhibited significant inhibitory effects on the hyphal growth and spore germination of Foc TR4, as well as severe destructive effects on its cell morphology and ultrastructure, and broad-spectrum antifungal activity. We also discovered that the inoculation of strain SCA4-21 significantly inhibited the infection of Foc TR4 in banana seedling corms, reduced the incidence index of banana *Fusarium* wilt, promoted the growth of banana seedlings, and enhances beneficial microbes and metabolic pathways, suggesting that strain SCA4-21 is a promising biocontrol agent.

## INTRODUCTION

Banana is a major source of starch in tropical and subtropical developing countries (1). Banana *Fusarium* wilt caused by Foc TR4 is a devastating soil-borne fungal disease. Foc TR4 is susceptible to almost all banana cultivars (2). In the absence of living host tissues, Foc TR4 can survive in soil as chlamydospores for over 20 years. Once the environmental conditions become favorable, the pathogen invades the root system of the banana plant and rapidly spreads. Its proliferating mycelium and secretions obstruct the vascular system, inducing water stress that leads to wilting and eventual mortality of the plant (3).

The utilization of biocontrol agents for managing banana *Fusarium* wilt is regarded as a viable alternative approach owing to its efficacy and eco-friendliness (4).

Rhizosphere *Streptomyces* spp. are ideal candidates for biocontrol agents against soil-borne diseases due to their ability to actively colonize plant root systems and survive for a long time even under extreme condition such as low nutrition and water availability in the form of spores (5–8). They have been studied extensively as potential biocontrol agents against fungal phytopathogens such as *Magnaporthe oryzae*, *Colletotrichum fragariae*, and *Colletotrichum gloeosporioides* (9–11). They protect crops from diseases by synthesizing diverse bioactive secondary metabolites, competing with pathogens for nutrients by producing substances like siderophores, or producing a large number of extracellular enzymes related to fungal cell wall degradation (6,12–14). We also founded that *Streptomyces* spp. has potential application prospects in the treatment of banana *Fusarium* wilt caused by Foc TR4 (15–17).

In our previous study, strain SCA4-21 was isolated from wheat rhizosphere soil in a dry-hot valley and identified as a new species of *Streptomyces*, named after *Streptomyces luomodiensis* sp. nov.. *S. luomodiensis* SCA4-21 exhibited excellent antagonistic activity against nine phytopathogenic fungi including Foc TR4 in vitro assay (18). In present study, the extracts of stain SCA4-21 were extracted and purified, and the half maximal effective concentration (EC_50_) of the extracts on Foc TR4 was determined. The extracts were also tested for their broad-spectrum antifungal activity against other eight plant pathogenic fungi. Additionally, the impact of the extracts on mycelial growth, spore germination, hyphal and spore morphology, as well as the ultrastructure of Foc TR4 was observed. Furthermore, volatile organic compounds of strain SCA4-21 were identified using gas chromatography-mass spectrometry (GC-MS). In addition, the biocontrol potential of strain SCA4-21 against banana *Fusarium* wilt caused by Foc TR4 and its effects on the growth of banana seedlings was evaluated. The effect of stain SCA4-21 on soil microbial communities was also analyzed through high-throughput sequencing. These findings highlight the excellent potential of strain SCA4-21 as a source of biocontrol agents.

### Result

### Extraction and purification of strain SCA4-21 extracts

The ethanol extracts of strain SCA4-21 fermentation broth were concentrated and adsorbed onto macroporous resins, then eluted with methanol gradient solutions to separate the antifungal extracts. The results showed that the extracts eluted with 100% methanol exhibited the highest mycelial inhibition rate (71.57±0.08) against Foc TR4, followed by the extracts eluted with 70% (31.53±0.15), 60% (17.67±0.25) and 50% (9.39±0.17) methanol, respectively (Fig. 1A). In fact, as the concentration of methanol increased, the antifungal activity of the extracts significantly enhanced. The extracts with eluted with 100% methanol was selected for next study due to its strong antifungal activity.

**1.**
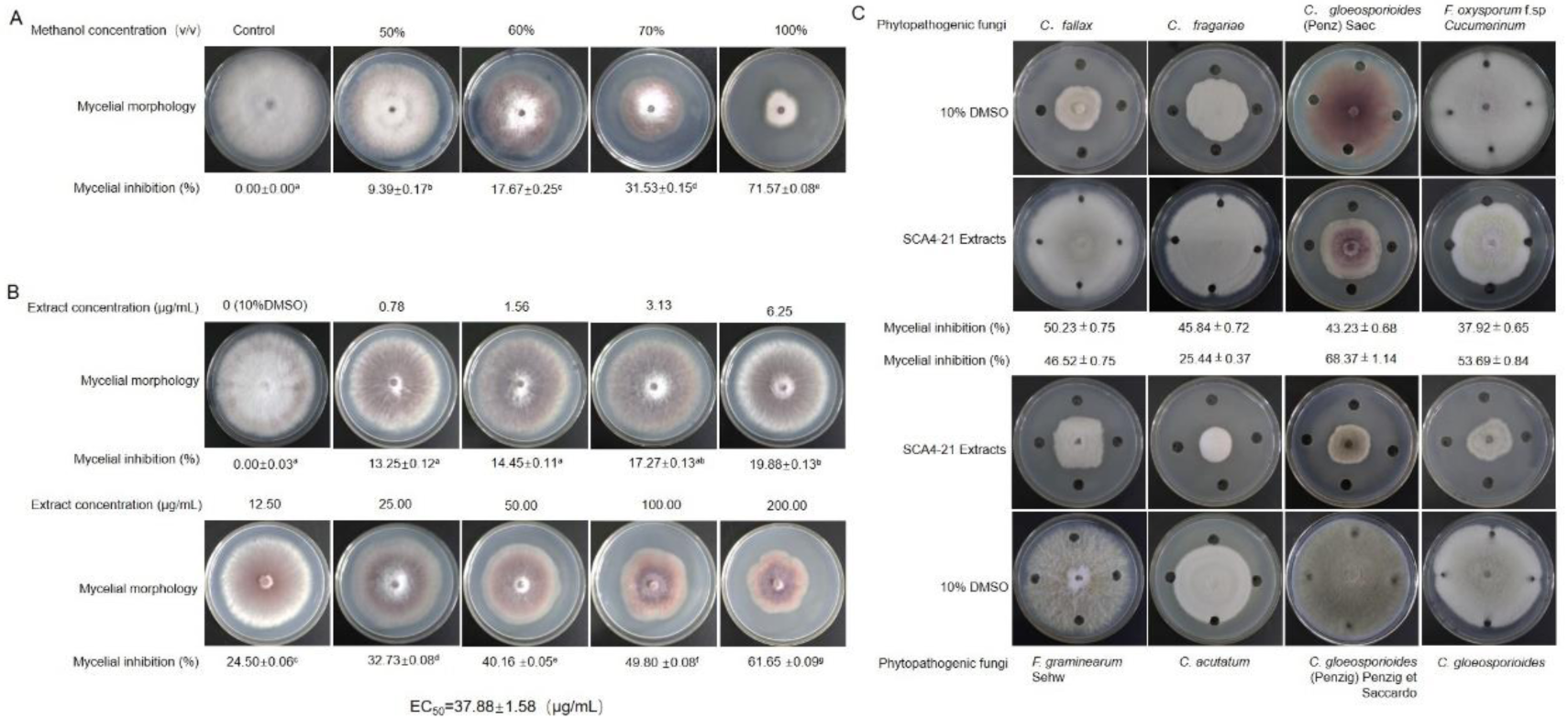
Isolated and activity assay of S. *luomodiensis* SCA4-21 extracts. (A) Effect of extracts separated by different gradient methanol solvents on the mycelial growth of Foc TR4. (B) Antifungal evaluation of extracts at different concentrations on the mycelial growth of Foc TR4. (C) Antifungal evaluation of extracts on the mycelial growth of other eight phytopathogenic fungi.

### Measuring the EC_50_ of strain SCA4-21 extracts on the mycelial growth of Foc TR4

The half maximal effective concentration (EC_50_) of strain SCA4-21 extracts on mycelial growth of Foc TR4 was determined. Mycelial inhibition showed a dose-dependent manner. After exposing Foc TR4 to the extracts at 28°C for 7 days, significant growth inhibition of the pathogen was observed in all concentration groups (0.78, 1.56, 3.13, 6.25, 12.5, 25, 50, 100, and 200 mg l^−1^). The inhibition of mycelial growth was as follows: 13.25±0.12, 14.45±0.11, 17.27±0.13, 19.88±0.13, 24.50±0.06, 32.73±0.08, 40.16 ±0.05, 49.80 ±0.08, 61.65 ±0.09, respectively. EC_50_ value is calculated as 37.88±1.58 µg mL−1 using toxicity regression equation (Fig. 1B).

### Strain SCA4-21 extracts exhibiting broad-spectrum antifungal activity

Strain SCA4-21 extracts showed effective antifungal activity against the tested strains of fungi. Compared to the control treated with 10% DMSO, the fungal hyphae treated with the extracts displayed significant inhibition. The inhibition rates for *C. gloeosporioides* (Penzig) Penzig et Saccardo, *C. gloeosporioides, C. gloeosporioides, F. graminearum Sehw, C. fragariae, C. gloeosporioides (Penz) Saec, F. oxysporum f.sp cucumerinum, C. acutatum* were 68.37%, 53.69%, 50.23%, 46.52%, 45.84%, 43.23%, 37.92 and 25.44, respectively (Fig. 1C).

### Strain SCA4-21 extracts significantly inhibiting spore germination of Foc TR4

The germination of spores was significantly inhibited after treated with extracts from strain SCA4-21(Fig. 2A). The inhibitory efficiency showed a positive correlation with the concentration of the extracts used in the treatment. Compared to the control group, the spore germination rates of Foc TR4 were 65%, 44% and 20% after treatment with 1 × EC_50_, 2× EC_50_, 4× EC_50_, respectively. The inhibition efficiencies were calculated as follows:15.6%, 42.9%, and 74.0%. Furthermore, almost complete inhibition of spore germination was observed when an extract concentration equivalent to 8 × EC_50_ was used (Fig. 2B).

**2.**
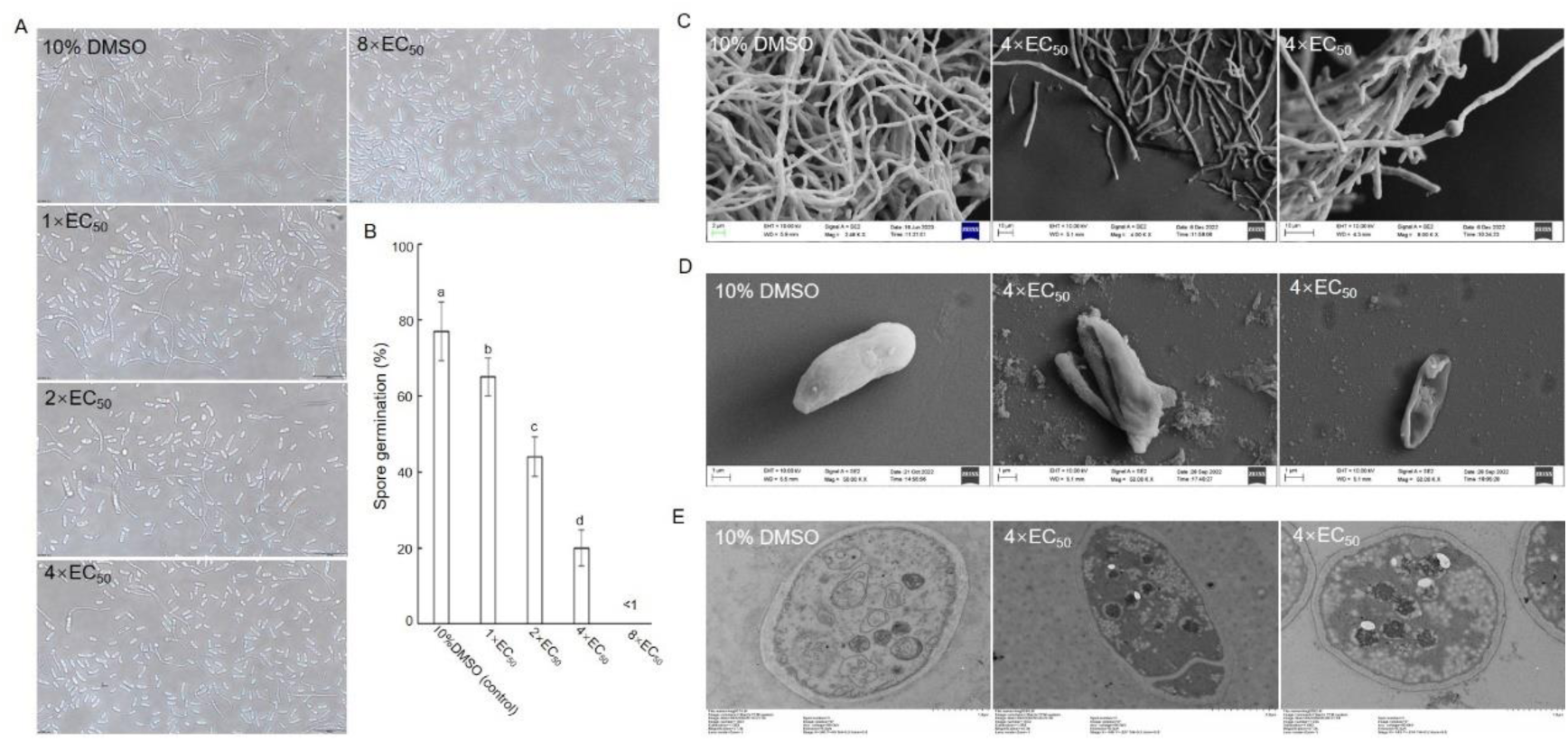
Effect of extracts on spore germination, spore and hyphal morphology as well as ultrastructure of Foc TR4 cells. (A) Inhibitory effects of extracts at different concentrations on the spore germination of Foc TR4. (B) Quantitative analysis of germination rate. Different letters represented a significant difference at the P < 0.05 level by the Duncan’s multiple range test. (C) Morphological characteristics of Foc TR4 hypha treated with 4 × EC_50_ of extracts. (D) Morphological characteristics of Foc TR4 spores treated with 4 × EC_50_ of extracts. (E) Ultrastructural characteristics of Foc TR4 cell treated with 4 × EC_50_ of extracts. The control groups were treated with 10% DMSO.

### Strain SCA4-21 extracts severely destroying morphology and ultrastructure of Foc TR4

SEM observation revealed that the control mycelium of Foc TR4 treated with 10% DMSO exhibited a smooth and intact morphology, while mycelium of Foc TR4 treated with the 4× EC_50_ extracts displayed notable alterations such as swelling and fusion (Fig. 2C). Simultaneously, the treated spores also exhibited a disrupted and concave morphology, whereas the control spores displayed a regular and planar appearance (Fig. 2D).

TEM analysis showed that the ultrastructure of Foc TR4 hyphae with the 4× EC_50_ extracts were also seriously damaged. Their organelles such as mitochondria and vacuoles disintegrated. The cytoplasm shrinks, the cell membrane becomes indistinct and the nucleus disappears. In contrast, the cell membrane and wall of the control group were clear and complete, Cell vacuoles are filled and mitochondrial cristae are clearly distinguishable (Fig. 2E).

### Identification of the volatile organic compounds of strain SCA4-21

GC–MS analysis was conducted to identify the volatile organic compounds of strain SCA4-21 that are likely responsible for its antifungal activities. by comparing the mass spectra of treatments with those of controls, 32 secondary metabolites of *S. luomodiensis* SCA4-21 were determined, including Oxetane, 2,3,4-trimethyl-, (2à,3 à,4á)-, Butanoic acid, methyl ester (Methyl butyrate), Ethene, fluoro-, Methyl isovalerate, Methyl tiglate, Pentanoic acid, 3-methyl-, methyl ester, Pentanoic acid, 4-methyl-, methyl ester, Hexanoic acid, methyl ester (methyl hexanoate), Thiopivalic acid, 6-Methyl-2-heptanone, 5-Methyl-2-heptanone, (E)-2-Hexenoic acid, methyl ester Methyl (E)-2-hexenoate, Furyl hydroxymethyl ketone, 3-Octanone Hexanoic acid, 5-methyl-, methyl ester, Heptanoic acid, methyl ester, (Fig. 3, Supplementary Fig. 1A-F and Table S1).

**3.**
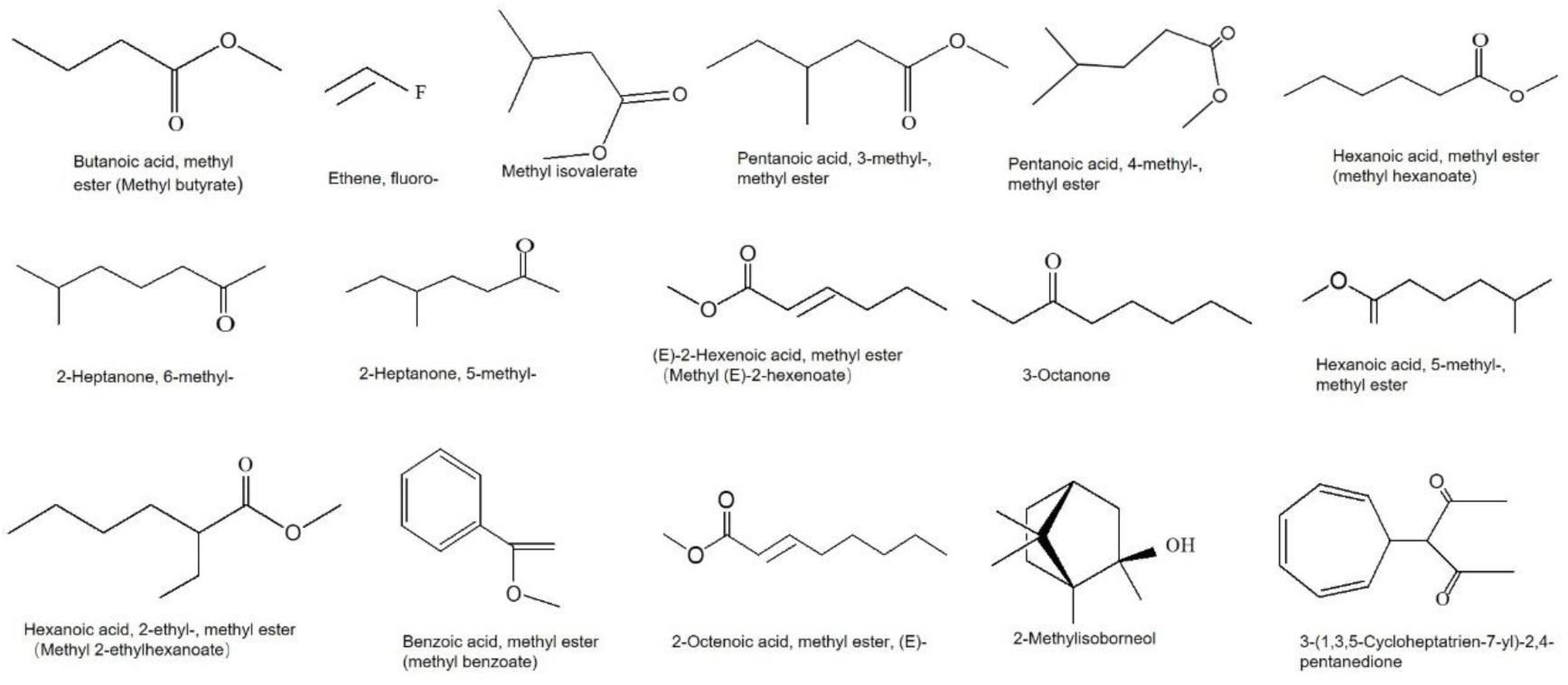
Chemical structures of the identified compounds of *S. luomodiensis* SCA4-21 by GC-MS.

### Strain SCA4-21 significantly reducing the incidence of banana *Fusarium* wilt disease

The pot experiment revealed that Strain SCA4-21 significantly reduced the incidence of banana wilt disease caused by Foc TR4 (Fig. 1A-B). Specifically, no evident chlorotic leaves were observed in negative control group (T1) where banana plantlets treated with sterilized SLM only. While, in positive control group (T2), the leaves of banana plantlets treated with sterilized SLM and GFP-Foc TR4 exhibited seriously chlorotic symptom, resulting in a disease index of 57.5%. In contrast, only a few old leaves of banana seedlings treated with GFP-Foc TR4+SCA4-21 (T3) showed yellow symptoms, leading to a significantly lower disease index of 23.4% compared to positive controls (Fig. 4A). The biocontrol efficiency of strain SCA4-21 against banana *Fusarium* wilt was 59.3% (Fig. 4B).

**4.**
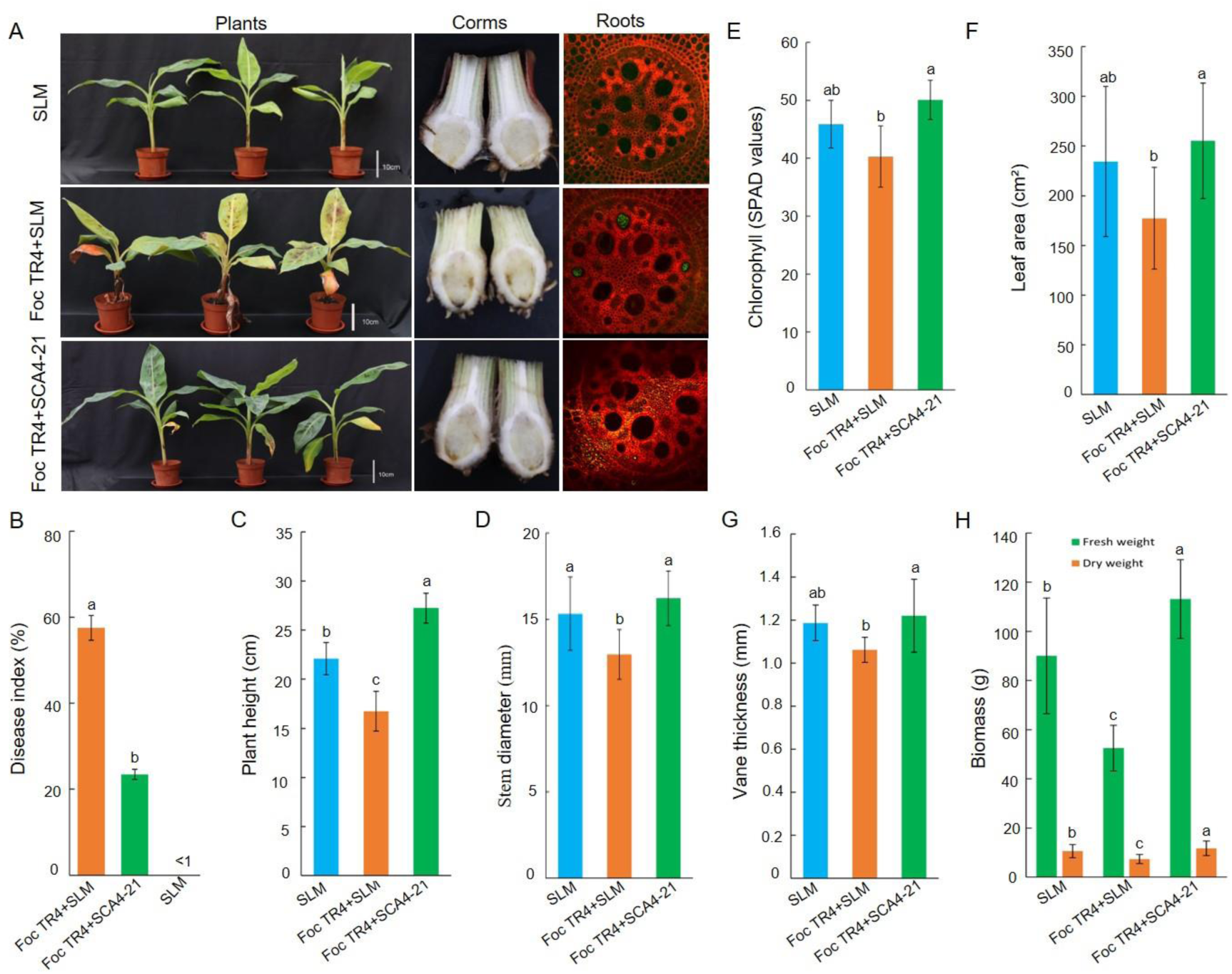
Biocontrol efficacy of *S. luomodiensis* SCA4-21 against banana *Fusarium* wilt and promotive effect on growth of banana seedlings. (A) In vivo effects of *S. luomodiensis* SCA4-21 on banana *Fusarium* wilt. (B) Quantitative analysis of disease index. Quantitative analysis of growth-related indices, including plant height (C), stem diameter (D), chlorophyll (E), leaf area (F), vane thickness (G), and biomass (H). Different lowercase letters indicated a significant difference among different treatments according to the Duncan’s multiple range test (p < 0.05).

Similarly, the corm sections of the T1, T2, and T3 groups displayed white, dark brown, and light brown colors respectively. Laser confocal microscope analysis further revealed that only a small amount of mycelium was observed in the outer cells of the T3 bulbs, unlike the corms in the T2 group where vascular cells were filled with Foc TR4 hyphae, while the corms in the T1 group were not infected by Foc TR4 (Fig. 4A).

### Strain SCA4-21 promoting significantly the growth of banana seedlings

The pot experiment also indicated that strain SCA4-21 significantly promoted the growth of banana seedlings (Fig. 4C-H). The T3 group, treated with SCA4-21+Foc TR4, exhibited significant increases in plant height (62.9%) (Fig. 4C), stem diameter (25.2%) (Fig. 4D), chlorophyll content (24.3%) (Fig. 4E), leaf area (43.9%) (Fig. 4F), vane thickness (14.9%) (Fig. 4G), fresh weight (115.5%) and dry weight (59.3) of banana seedlings 60 days after planting when compared to the T2 group treated with SLM+Foc TR4 (Fig. 4H).

Compared to the T1 group treated with SLM, the T3 group also showed significant increases in plant height (23.3%), fresh weight (25.6%), and dry weight (9.6%). Additionally, stem diameter, chlorophyll content, leaf area, and leaf thickness increased by 5.9%, 9.1%, 8.9%, and 2.8% respectively; however, there was no statistically significant difference between these four indicators in the two groups.

### Strain SCA4-21 shaping structure of banana rhizosphere bacterial community

720,187 pairs of raw reads were obtained from 9 soil samples by sequencing the V3+V4 regions of bacterial 16S rRNA. These bacterial raw reads were submitted to NCBI’s Sequence Read Archive database (accession number: SRR26906769-74, SRR26906783-4). Subsequently, after filtering, cutting and splicing, a total of 718,594 pairs of clean reads were obtained (Table 1). After the chimera was removed, the obtained 482,135 non-chimeric clean reads were clustered into 11210 distinct bacterial Operational Taxonomic Units (OTUs) with 97% similarity. Among them, 1732, 1899 and 1862 OTUs were derived fromT1 group soil samples treated with sterilized SLM. while 1946, 1863 and 1851 OTUs were derived from T2 group soil samples treated with sterilized SLM and GFP-Foc TR4. Similarly, 1602, 1852 and 1647 OTUs were derived from T3 group soil samples treated with Foc TR4+SCA4-21(Fig. 5A). Venn analysis revealed that T1, T2, and T3 groups had 3201,3408 and 3145 unique bacteria OTUs respectively (Fig. 5B).

**Table 1.** Statistics of bacterial 16S rRNA gene sequencing data processing results for soil samples with different treatments.

| Sample ID | Raw Reads | Clean Reads | Denoised Reads | Merged Reads | Non-chimeric Reads |
| --- | --- | --- | --- | --- | --- |
| SLM | 79966 | 79771 | 74069 | 59601 | 50140 |
| SLM | 80121 | 79953 | 74192 | 58275 | 47570 |
| SLM | 79926 | 79767 | 74365 | 60998 | 51339 |
| Foc TR4+SLM | 80039 | 79878 | 74308 | 66628 | 59763 |
| Foc TR4+SLM | 80123 | 79937 | 73295 | 60729 | 51707 |
| Foc TR4+SLM | 79909 | 79735 | 72242 | 59815 | 51019 |
| Foc TR4+SCA4-21 | 80362 | 80187 | 74852 | 68004 | 58432 |
| Foc TR4+SCA4-21 | 79863 | 79684 | 74327 | 70695 | 62599 |
| Foc TR4+SCA4-21 | 79878 | 79682 | 73340 | 62188 | 49566 |
| Total | 720187 | 718594 | 664990 | 566933 | 482135 |

**5.**
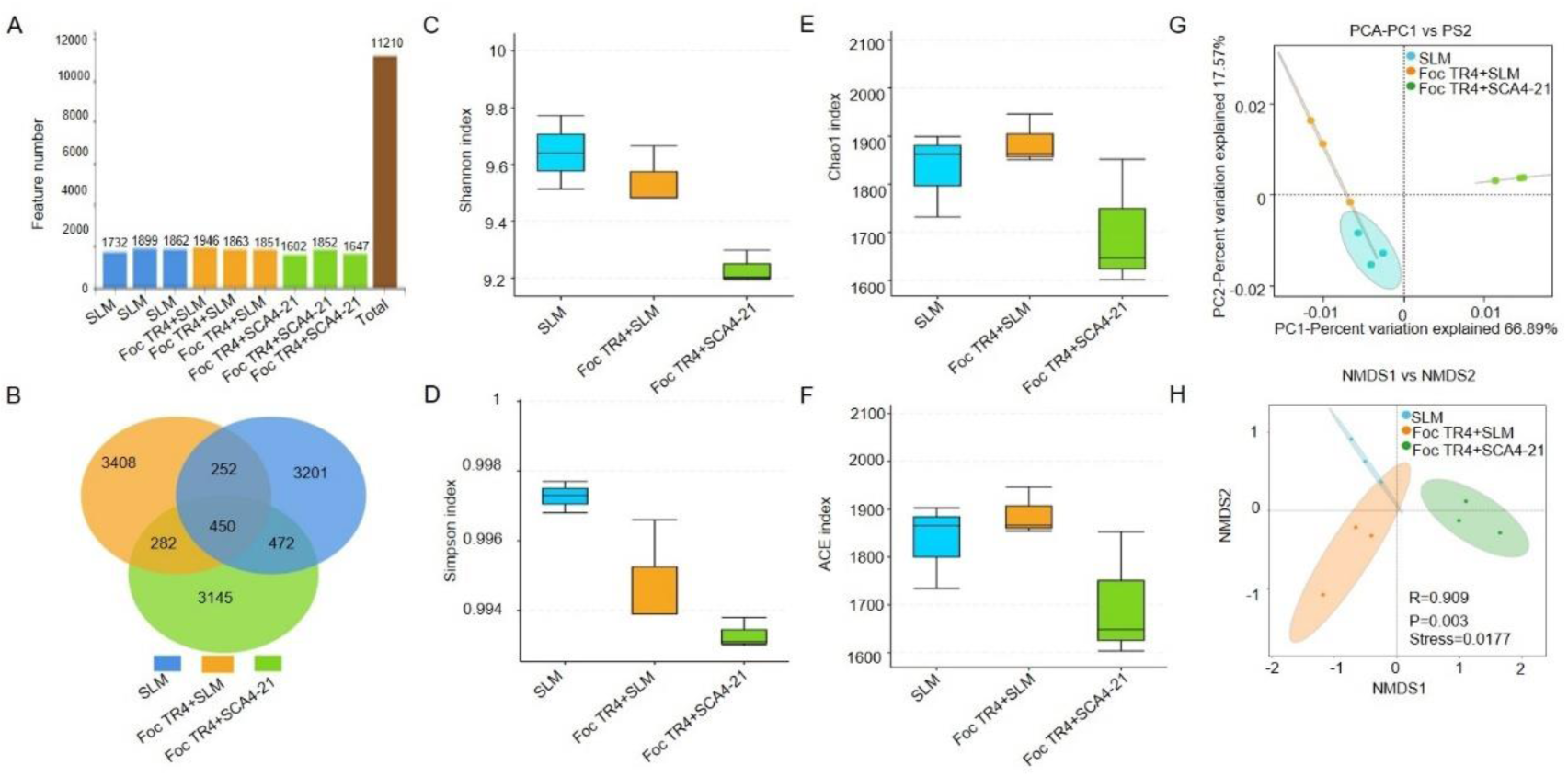
Effect of S. *luomodiensis* SCA4-21 on diversity of rhizosphere soil bacterial communities. (A) Identification of OTUs in different soil samples. (B) Venn diagram of different treatment. (C, D) Diversity indices Shannon and Simpson. (E, F) Richness indices Chao1 and ACE. (G) Principal component analysis of different treatment. (H) Nonmetric multidimensional scaling analysis of different treatment.

Analysis of bacterial alpha diversity showed that the diversity indices Shannon and Simpson in T1 group soils were higher than those in T2 and T3 group soils (Fig. 5C-D). While, T2 group soils showed higher the richness indices ACE and Chao1, followed by the T1 and T3 group soils (Fig. 5E-F). However, there were no statistically significant differences observed in the bacterial the diversity indices Shannon and Simpson, as well as the richness indices ACE and Chao1, among the three treatments (Fig. 5C-F and Supplementary Table S2).

Principal component analysis (PCA) revealed a tendency for different samples of the same treatment to cluster together, indicating a relatively high similarity in bacterial community composition among samples within the same group. PCA results also showed that PC1 explained 66.89% of the total variation, PC2 explained 17.98% of the total variation, suggesting good separation in bacterial community composition among the treatment groups (Fig. 5G). Nonmetric multidimensional scaling (NMDS) analysis result clearly showed significant variations in bacterial community composition across the different treatments (ANOSIM, R = 0.909, P=0.003), and low stress value (stress=0.0017<0.05) indicating that the analysis results have excellent representativeness (Fig. 5H).

Taxonomic annotations showed that a total of 11,210 bacterial OTUs were classified into 2 kingdoms, 40 phyla, 99 classes, 265 orders, 515 families, 971 genera and 1142 species (Table 2). The top 20 bacterial genera in terms of relative abundance were displayed using bar chart (Fig. 6A) and heatmap (Supplementary Fig. S2A), including unclassified-*Xanthobacteraceae* (5.32%, 4.97% and 2.29%, respectively), *Bacillus* (0.30%, 0, and 9.55% respectively) unclassified-*Acidobacteriales* (5.49%, 3.54%, and 2.19% respectively), unclassified-*Bacteria* (3.17%, 3.37%, and 1.77% respectively), unclassified-*Cyanobacteriales* (1.06%, 5.16%, and 0.32% respectively), *Burkholderia*-*Caballeronia*-*Paraburkholderia* (1.81%, 0.91%, and 3.75% respectively), unclassified-*Elsterales* (3.37%, 2.18%, and 0.85% respectively), *Haliangium* (2.53%, 2.43%, and 1.34% respectively), *Pantoea* (1.09%, 4.74%, and 0.05% respectively), *Bryobacter* (3.22%, 1.38%, and 0.96% respectively), unclassified-*Gemmatimonadaceae* (1.94%, 2.30%, and 1.31% respectively), *Sphingomonas* (0.91%, 0.82%, and 2.95% respectively), *Cupriavidus* (0.62%, 0.48%, and 3.37% respectively), unclassified-*Comamonadaceae* (1.18%, 2.12%, and 1.14% respectively), unclassified-SBR1031(2.49%, 1.52%, and 0.25% respectively), unclassified-*Alphaproteobacteria* (1.86%, 1.73%, and 0.63% respectively), *Massilia* (0.18%, 0.05%, and 3.98% respectively), *Pseudomonas* (0.33, 0.12%, and 3.71% respectively), uncultured-*Acidobacteria*-*bacterium* (1.58%, 1.74%, and 0.59% respectively), *Streptomyces*(0.48%, 0.31%, and 2.30% respectively) (Supplementary Table S3).

**Table 2.**
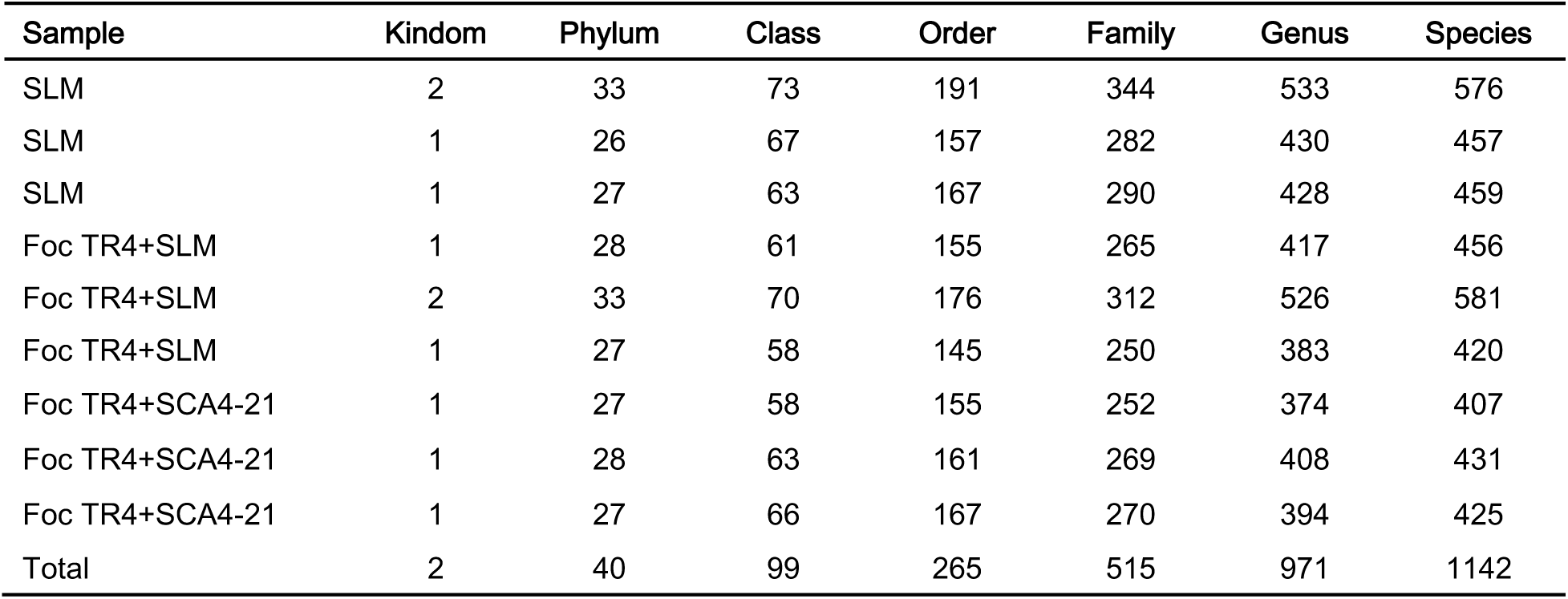
Bacterial species identified at different levels in soil samples under different treatment.

| Sample | Kindom | Phylum | Class | Order | Family | Genus | Species |
| --- | --- | --- | --- | --- | --- | --- | --- |
| SLM | 2 | 33 | 73 | 191 | 344 | 533 | 576 |
| SLM | 1 | 26 | 67 | 157 | 282 | 430 | 457 |
| SLM | 1 | 27 | 63 | 167 | 290 | 428 | 459 |
| Foc TR4+SLM | 1 | 28 | 61 | 155 | 265 | 417 | 456 |
| Foc TR4+SLM | 2 | 33 | 70 | 176 | 312 | 526 | 581 |
| Foc TR4+SLM | 1 | 27 | 58 | 145 | 250 | 383 | 420 |
| Foc TR4+SCA4-21 | 1 | 27 | 58 | 155 | 252 | 374 | 407 |
| Foc TR4+SCA4-21 | 1 | 28 | 63 | 161 | 269 | 408 | 431 |
| Foc TR4+SCA4-21 | 1 | 27 | 66 | 167 | 270 | 394 | 425 |
| Total | 2 | 40 | 99 | 265 | 515 | 971 | 1142 |

**6.**
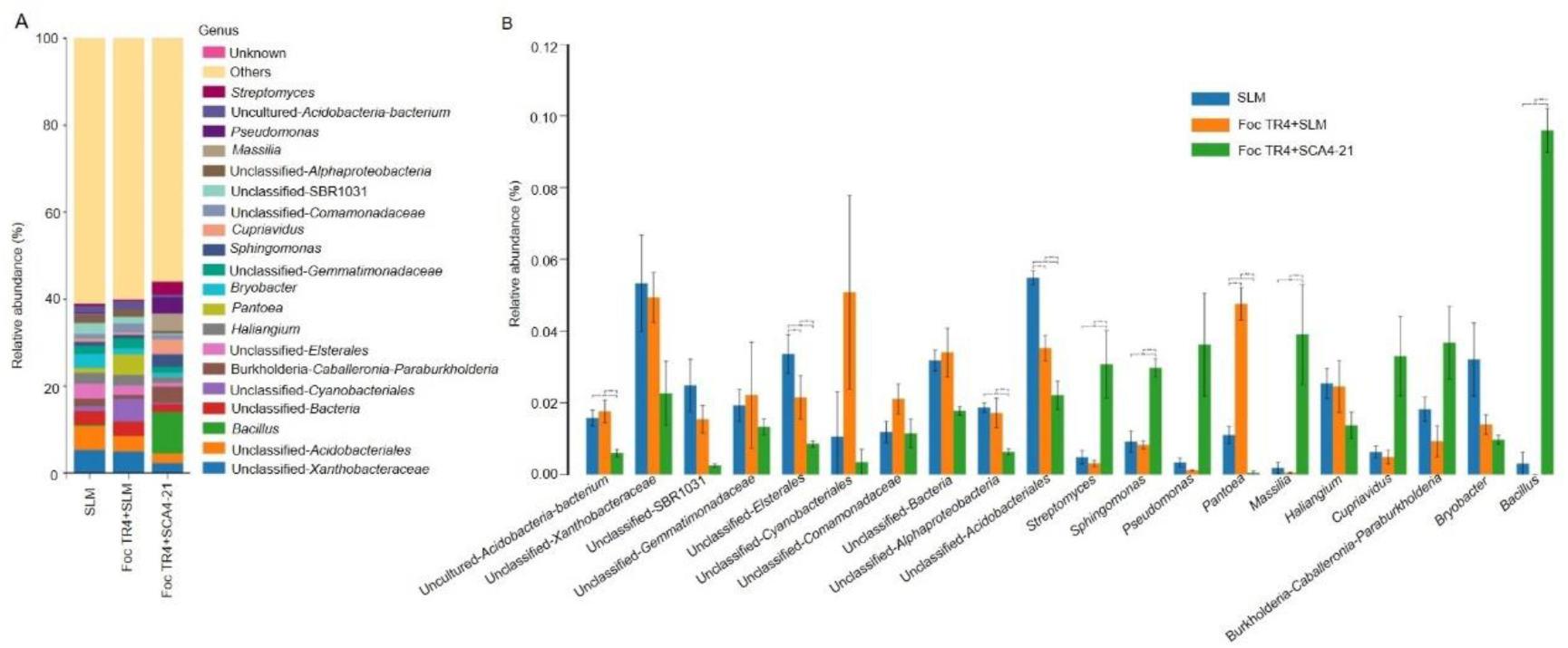
Effect of S. *luomodiensis* SCA4-21 on composition of rhizosphere soil bacterial communities. (A) Relative abundances of top 20 bacterial genus under each treatment. (B) Analysis of intergroup differences in the abundance of the top 20 bacterial genera using the analysis of variance (ANOVA) method.

ANOVA method was used to analyze the differences among the top 20 bacterial genera in three treatments (Fig. 6B). Compared to T1 and T2 groups soils, the soil samples of the T3 group treated with Foc TR4+SCA4-21 exhibited significantly higher abundances of *Streptomyces, Bacillus*, *Sphingomonas*, and *Massilia*. Conversely, the abundances of *uncultured-Acidobacteria-bacterium, unclassified-Elsterales, unclassified-Alphaproteobacteria,* unclassified-*Acidobacteriales* and *Pantoea* is significantly lower in T3 group soils than in T1 and T2 group soils.

Bacterial functional profiles based on the Kyoto Encyclopedia of Genes and Genomes (KEGG) was predicted via Tax4Fun (Fig. 7A-C). Compared to groups T1 and T2, T3 group exhibited significant enrichment in pathways related to carbohydrate metabolism, amino acid metabolism, lipid metabolism, metabolism of terpenoids and polyketides, and biosynthesis of other secondary metabolites (Fig. 7A-B). Conversely, when comparing T1 and T3 groups, pathways such as nucleotide metabolism and translation were significantly increased in group T2 (Fig. 7A, 7C). Additionally, pathways such as folding, sorting and degradation, and replication and repair were significantly increased in both T2 and T3 groups compared to group T1 (Fig. 7B-C). In order to explore the competitive or cooperative relationships between *Streptomyces* and other bacterial genera in soil samples treated differently, Spearman’s rank correlation coefficients were calculated and co-occurrence network of top 20 bacterial genera by abundance was constructed. The results revealed that the relative abundance of *streptomyces* was positive with *Sphingomonas*, *Burkholderia*-*Caballeronia*-*Paraburkholderia*, *Massilia, Pseudomonas, Bacillus,* and *Cupriavidus*, while it was negatively correlated with *Pantoea* and unclassified *Cyanobacteriales* (Fig. 7D).

**7.**
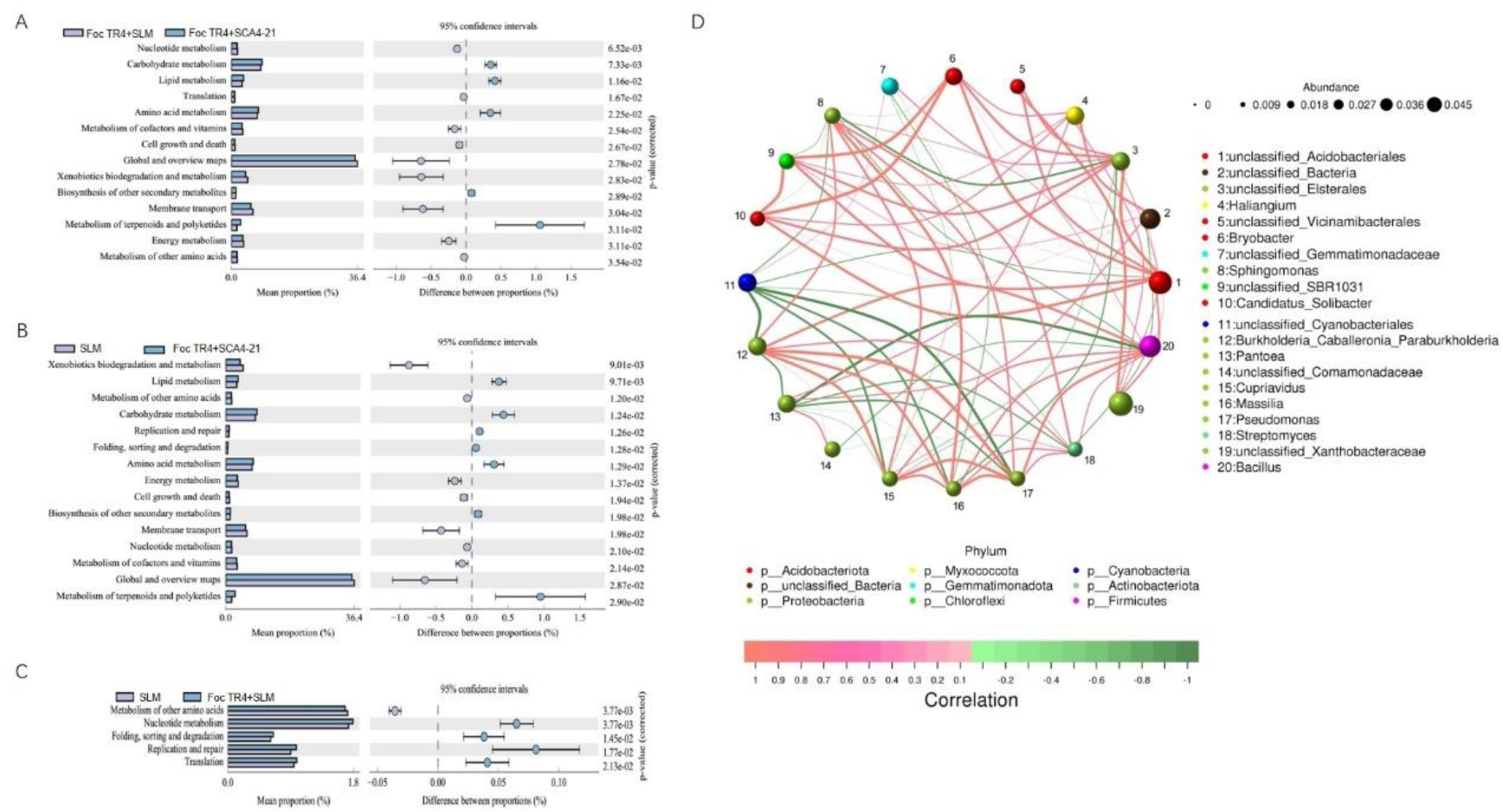
Effect of S. *luomodiensis* SCA4-21 on function and correlation of rhizosphere soil bacteria. (A) Functional profile prediction of bacteria community using Tax4Fun2. (B) Network correlation analysis of top 20 bacterial genus.

### Strain SCA4-21 monitoring structure of banana rhizosphere fungal community

Fungal ITS analysis of 9 soil samples yielded 720,798 pairs of raw reads. These fungal raw reads were submitted to NCBI’s Sequence Read Archive database (accession number: SRR26906767, SRR26906775-82). Following splicing, chimaera removal, and denoising procedures, we obtained a set of high-quality clean reads comprising 690,176 pairs (Table 3). These authentic biological sequences were subsequently clustered into 2454 fungal OTUs at a similarity threshold of 97%. Specifically, the three soil samples from T1 group contributed 309, 411 and 590 OTUs, respectively. The three soil samples from T2 group comprised 316, 446 and 429 OTUs, respectively. The three soil samples from T3 group generated 413, 341 and 480 OTUs, respectively (Fig. 8A). Venn analysis demonstrated that T1, T2, and T3 groups had 677,615 and 719 unique fungal OTUs respectively (Fig. 8B).

**8.**
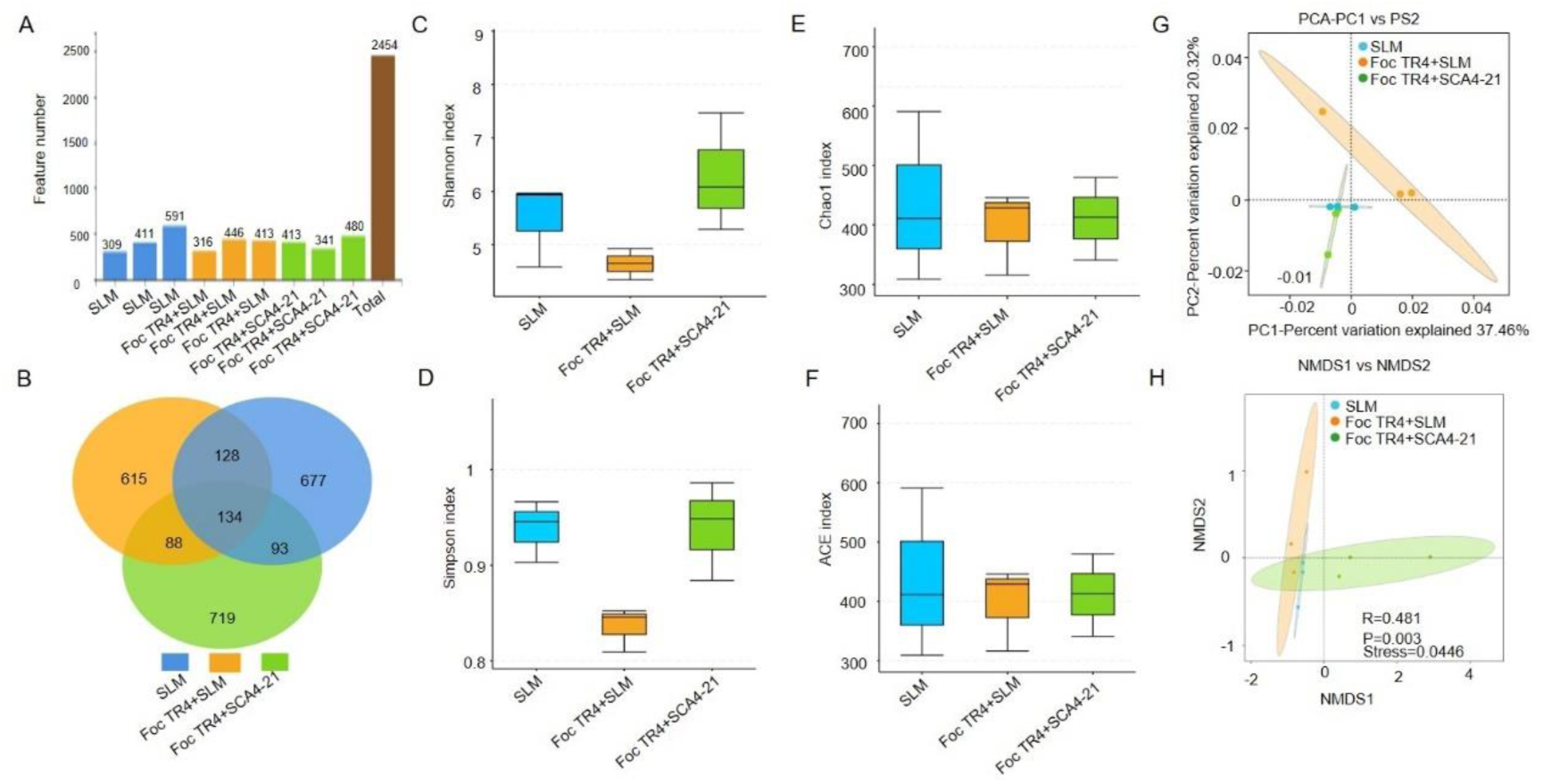
Effect of S. *luomodiensis* SCA4-21 on diversity of rhizosphere soil fungal communities. (A) Identification of OTUs in different soil samples. (B) Venn diagram of different treatment. (C, D) Diversity indices Shannon and Simpson. (E, F) Richness indices Chao1 and ACE. (G) Principal component analysis of different treatment. (H) Nonmetric multidimensional scaling analysis of different treatment.

**Table 3.** Statistics of fungal internal transcribed spacer sequencing data processing results for soil samples with different treatments.

| Sample ID | Raw Reads | Clean Reads | Denoised Reads | Merged Reads | Non-chimeric Reads |
| --- | --- | --- | --- | --- | --- |
| SLM | 79906 | 79566 | 78867 | 77548 | 75255 |
| SLM | 79890 | 79505 | 78644 | 77148 | 77023 |
| SLM | 80044 | 79637 | 78167 | 76089 | 74962 |
| Foc TR4+SLM | 80085 | 79814 | 79456 | 78027 | 77692 |
| Foc TR4+SLM | 80424 | 80067 | 79277 | 77274 | 77216 |
| Foc TR4+SLM | 80110 | 79831 | 79108 | 76998 | 76793 |
| Foc TR4+SCA4-21 | 80324 | 79973 | 79605 | 77926 | 77814 |
| Foc TR4+SCA4-21 | 79984 | 79581 | 79301 | 78148 | 77432 |
| Foc TR4+SCA4-21 | 80031 | 79698 | 79056 | 76523 | 75989 |
| Total | 720798 | 717672 | 711481 | 695681 | 690176 |

**Table 4.** Fungal species identified at different levels in soil samples under different treatments.

| Sample | Kindom | Phylum | Class | Order | Family | Genus | Species |
| --- | --- | --- | --- | --- | --- | --- | --- |
| SLM | 1 | 9 | 24 | 53 | 85 | 106 | 111 |
| SLM | 1 | 8 | 26 | 59 | 93 | 133 | 144 |
| SLM | 1 | 10 | 32 | 72 | 132 | 200 | 227 |
| Foc TR4+SLM | 1 | 8 | 24 | 59 | 92 | 131 | 142 |
| Foc TR4+SLM | 1 | 10 | 29 | 69 | 111 | 139 | 142 |
| Foc TR4+SLM | 1 | 10 | 27 | 61 | 97 | 129 | 136 |
| Foc TR4+SCA4-21 | 1 | 11 | 30 | 70 | 110 | 166 | 196 |
| Foc TR4+SCA4-21 | 1 | 9 | 29 | 54 | 98 | 140 | 160 |
| Foc TR4+SCA4-21 | 1 | 10 | 26 | 65 | 120 | 173 | 189 |
| Total | 1 | 11 | 42 | 105 | 217 | 400 | 537 |

Analysis of fungal alpha diversity indicated that the diversity indices Shannon and Simpson in T3 group soils were higher than those in T1 and T2 group soils (Fig. 8C-D), while the richness indices Chao1 and ACE are slightly higher in T2 group soils than in T1 and T3 group soils (Fig. 8E-F). However, no statistically significant differences were observed in the fungal diversity indices Shannon and Simpson, as well as the richness indices ACE and Chao1, among the three treatments (Fig. 8C-F and Supplementary Table S4).

PCA results revealed that the fungal community composition in group T2 differs significantly from groups T1 and T3, while the difference in fungal community composition between groups T1 and T3 is relatively small (Fig. 8G). Furthermore, the PCA results showed that the T2 group had a positive contribution to the total variation, while the T1 and T3 groups had a negative contribution (Fig. 8G). The PCA analysis also revealed that PC1 accounted for 37.46% of the total variation, and PC2 explained 20.32% of the total variation, indicating a clear separation in fungal community composition among the treatment groups (Fig. 8G). The NMDS analysis revealed significant differences in fungal community composition among the different treatments (ANOSIM, R = 0.481, P = 0.003) with a low stress value (stress = 0.0446 < 0.05), suggesting the highly representative nature of the analysis results (Fig. 8H). Taxonomic annotations showed that a total of 2454 fungal OTUs were identified into 1 kingdom, 11 phyla, 42 classes, 105 orders, 217 families, 400 genera and 537 species (Table 3). The fungal genera in the top 20% relative abundance were illustrated using bar charts (Fig. 9A) and heatmaps (Supplementary Fig. S2B). These identified dominant genera included *Fusarium* (6.65%, 37.29% and 16,92%, respectively), unclassified-*Sordariomycetes* (7.34%,21.36% and 6.98%, respectively), unclassified-*Basidiomycota* (15.13%,3.68% and 12.06%, respectively), unclassified-*Ascomycota* (7.56%, 6.78% and 3.89%, respectively), unclassified-*Agaricomycetes* (8.96%, 6.35% and 1.58%, respectively), unclassified-Fungi (4.59%, 3.35% and 3.77%, respectively), unclassified-*Dothideomycetes* (1.52%, 4.03% and 1.17%, respectively), *Gymnopilus* (5.56%, 0.01% and 0, respectively), *Conocybe* (2.43%, 0.65% and 2.48%, respectively), *Gibellulopsis*(0.39%, 0.3% and 2.51%, respectively), *Xenomyrothecium* (0.12%,0.16% and 2.74%, respectively), unclassified-*Thelephoraceae* (2.67%, 0 and 0, respectively), *Purpureocillium* (0.34%, 0 and 2.28%, respectively), *Lycoperdon* (1.3%, 0 and 1.28%, respectively) *Mortierella* (0.23%, 0.06% and 2.00%, respectively), *Cladosporium* (0.52%, 0.33% and 1.41%, respectively), unclassified-*Glomeraceae* (0.86%, 0.68% and 0.33%, respectively), *Rhizophagus*(1.11%, 0.54% and 0.16%, respectively), *Arthrobotrys* (0.05%, 0.17% and 1.27%, respectively) and unclassified-*Glomeromycota* (0.56%, 0.54% and 0.34%, respectively) (Supplementary Table S5).

**9.**
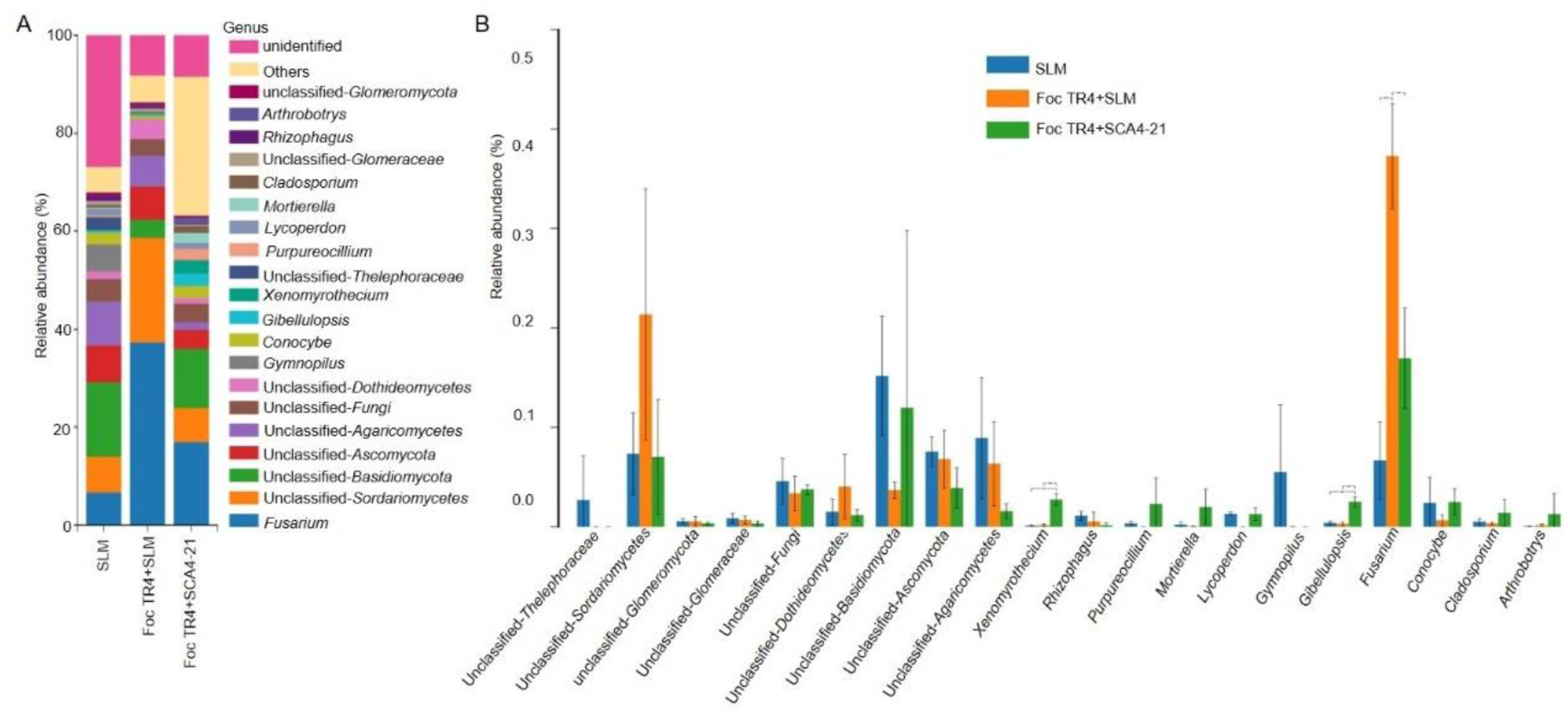
Effect of S. *luomodiensis* SCA4-21 on composition of rhizosphere soil fungal communities. (A) Relative abundances of top 20 bacterial genus under each treatment. (B) Analysis of intergroup differences in the abundance of the top 20 bacterial genera using the analysis of variance (ANOVA) method.

ANOVA (Analysis of Variance) method was also used to analyze the differences among the top 20 fungal genera in three treatments (Fig. 9B). Compared to Compared to T1 and T2 groups soils, the T3 groups soil showed significantly higher abundances of *Gibellulopsis* and *Xenomyrothecium.* In contrast, the abundances of *Fusarium* significantly decreased in T3 groups soil than that in T2 groups soil. Although the *Fusarium* abundance in T3 group soil was still higher than that in T1 group soil, there was no significant difference between them.

## Discussion

*Streptomyces* spp. represents a significant source of microbial bioagents. The production of antimicrobial bioactive substances undeniably constitutes a crucial mechanism through which *Streptomyces* spp. exert their function as biocontrol agents. Behera et al. demonstrated that *S. chilikensis* RC1830 exhibited in vitro antifungal activity against *F. oxysporum* through the production of metabolites, leading to a significant reduction (80.51%) in disease severity index of rice wilt disease in pot experiments (19). Chen et al. also reported that *S. plicatus* B4-7 showed the ability to synthesize the antifungal compound borrelidin, resulting in a significant reduction of 75% in crown rot disease incidence (20). Our previous studies have also shown that some *Streptomyces* strains such as *Streptomyces* sp. CB-75 and *Streptomyces* sp. YYS-7 significantly decreased the incidence of *Fusarium* wilt of banana by inhibiting mycelium growth, spore germination and destroying cell ultrastructure of Foc TR4 through secondary metabolites (15, 21).

In this study, we also discovered that extracts of *S. luomodiensis* SCA4-21 significantly inhibited the growth of Foc TR4 hyphae (Fig. 1A-B) and spore germination (Fig. 2A-B), severely disrupted the ultrastructure of Foc TR4 hyphal cells and spore morphology (Fig. 2C-E). The extracts also showed broad-spectrum antifungal activity against eight other phytopathogenic fungi (Fig. 1C). GC-MS analysis revealed that strain SCA4-21 produces a total of 32 volatile organic compounds, including Butanoic acid, methyl ester, Methyl isovalerate, 5-Methyl-2-heptanone, 6-Methyl-2-heptanone, 3-Octanone, Hexanoic acid, 2-ethyl-, methyl ester, and Benzoic acid, methyl ester, among others (Fig. 3 and Supplementary Table S5). The previous studies have indicated that these compounds possess specific biological activities. Khan et al. reported that Butanoic acid, methyl ester (Methyl butyrate), at a concentration as low as 0.01 M, induced significant cytotoxicity in the human breast cancer cell line MDA-MB-231 (22). Ayaz et al. discovered that Methyl isovalerate exhibited strong nematicidal activity, achieving a mortality rate of 83% (23). Morita et al. and Zhang et al. demonstrated that 5-Methyl-2-heptanone and 6-Methyl-2-heptanone exhibit potent antifungal activities against *Penicillium italicum* and *Alternaria solani*, respectively (24–25). Dotson et al. reported that the *Streptomyces* volatile 3-octanone could modulate auxin/cytokinin levels, promoting growth in *Arabidopsis thaliana* via the gene family *KISS ME DEADLY* (26). Our previous findings also revealed that Hexanoic acid, 2-ethyl-, methyl ester Methyl 2-ethylhexanoate, derived from *Streptomyces corchorusii* CG-G2, exhibits significant antifungal activity against *Colletotrichum gloeosporioides* (27). Lima et al. demonstrated that Benzoic acid, methyl ester (methyl benzoate) exhibits antifungal activity against *Candida albicans* (28). These results suggest that the compounds identified are responsible for the antifungal activity of S. *luomodiensis* SCA4-21. In this study, we also evaluated biocontrol efficiency of strain SCA4-21 against banana *Fusarium* wilt. Our results demonstrated that strain SCA4-21 could suppress mycelial infection of Foc TR4 in roots, resulting in a remarkable reduction by 59.3% in disease incidence banana *Fusarium* wilt, and promote significantly the growth of banana seedlings (Fig. 4A-H). The rhizosphere microbiome serves as the first line of defense against soilborne pathogens (29). Therefore, it is essential to analyze the effects of bioagent inoculation on the rhizosphere microbiome in the biocontrol of soil-borne diseases. Our results of current high-throughput sequencing showed a significant increase in the abundances of the genus *Streptomyces, Bacillus, Sphingomonas*, and *Massilia* in the T3 group (treated with Foc TR4+SCA4-21), compared to both the T1 group (treated with sterilized SLM) and the T2 group (treated with sterilized SLM+ Foc TR4) (Fig. 6A-B). Moreover, T3 group also exhibited higher the abundances of the genus *Pseudomonas, Burkholderia-Caballeronia-Paraburkholderia* and *Cupriavidus* than the other two groups (Fig. 6A-B). We also observed that the abundance of the bacterial genus *Pantoea* significantly decreased in the T3 group compared to that in the T1 and T2 groups (Fig. 6A-B). Therefore, we propose that the introduction of strain SCA4-21 may directly or in directly attribute to the increased abundance of *Streptomyces*, *Bacillus*, *Sphingomonas*, *Massilia*, *Burkholderia*-*Caballeronia*-*Paraburkholderia*, and *Cupriavidus*, while simultaneously reducing the prevalence of *Pantoea* in the T3 group. Spearman network correlation analysis was conducted to identify associations between *Streptomyces* and other bacterial genera, revealing a positive correlation between *Streptomyces* and *Bacillus*, *Sphingomonas*, *Massilia*, *Burkholderia*-*Caballeronia*-*Paraburkholderia*, and *Cupriavidus*, while showing a negative correlation with *Pantoea* genus (Fig. 7D). This further confirms the regulatory effect of strain SCA4-21 on these bacteria.

The accumulated data showed that the genera *Bacillus*, *Sphingomona* and *Pseudomonas* are important resources for biological control agents (30–34). Actually, these beneficial bacteria are often recruited and enriched in plant roots by microbial bioagents, playing an important role in defending against soil-borne diseases. Zhang et al. reported that *Trichoderma asperellum* M45a triggered watermelon resistance to *Fusarium* wilt by regulating rhizosphere microbes, including increasing the abundance of the genera *pseudomonas* and *Sphingomonas* in soil (35). Yang et al. found that *Streptomyces aureoverticillatus* HN6 could colonize the banana rhizosphere soil and effectively inhibited banana *Fusarium* wilt disease by attracting *Bacillus* and *Pseudomonas* to colonize the banana plant rhizosphere and decreasing the relative abundances of *Fusarium* (36). Our previous studies also revealed that the genera *Bacillus*, *Sphingomonas*, and *Pseudomonas* were significantly enriched in the rhizosphere soil of banana plants free from *Fusarium* wilt compared to those infected by the fungus (37). Additionally, the genus *Massilia* has been frequently documented as a colonizer of both the rhizosphere and endorhizal environments (Ofek et al., 2012; Kuffner et al., 2010). Some *Massilia* isolates have also been found to control plant pathogens by secreting siderophore or extracellular enzymes (Hrynkiewicz et al., 2010; Grönemeyer et al., 2011). The applications of the genera *Burkholderia*-*Caballeronia*-*Paraburkholderia* and *Cupriavidus* in biocontrol remain undocumented; however, they are frequently identified as dominant genera in various foods, such as broad-bean paste and tea, suggesting their role as beneficial bacteria (36; Zeng et al., 2020). However, *Pantoea* spp. are frequently recognized as plant pathogens (Wensing et al., 2010; Dussault et al., 2017). Mwaheb et al. (2017) also demonstrated that the inoculation of *Hirsutella minnesotensis* significantly reduced the abundance of the genus *Pantoea* in nematode-infested soils.

Our findings also indicate that the introduction of Foc TR4 significantly enhanced the abundance of the fungal genus *Fusarium* in the T2 group, while the inoculation of strain SCA4-21 significantly decreased *Fusarium* abundance in the T3 group (Fig. 9A-B). This suggests that strain SCA4-21 inhibits the proliferation of Foc TR4 in banana rhizosphere soil. The severity of banana *Fusarium* wilt is accompanied by an increase in *Fusarium* colonization in the field (37). The microbial bioagents for banana *Fusarium* wilt often exhibit an inhibition of the proliferation of *Fusarium* in soil. Shen et al. revealed that *Bacillus amyloliquefaciens* NJN-6 could more effectively control *Fusarium* wilt disease by suppressing *Fusarium* growth in the field (38). We also observed that the relative abundances of the fungal genera *Mortierella*, *Purpureocillium, Gibellulopsis* and *Xenomyrothecium* in the T3 group soils were significantly higher than those in the T1 and T2 group soils (Fig. 9A-B), Especially the abundance of *Mortierella* is approximately 9 and 33 times higher in the T3 group than in the T1 and T2 groups, respectively (Supplementary Table S5), suggesting strain SCA4-21 greatly promoted the enrichment of these fungal genus in the banana rhizosphere soil. *Mortierella* spp. are frequently recognized as beneficial fungi. Soman et al. found that *Mortierella vinacea* exhibited antifungal and antibacterial activity (39). Li et al. (2014) reported that relative abundance of *Mortierella* dropped significantly in the consecutive peanut monoculturing fields. Ye et al. found that an increase in the relative abundance of *Fusarium* leads to a decrease in the relative abundance of *Mortierella* (40). We previously indicated that *Mortierella* (36.64%) was predominant fungal genus in the *Fusarium* wilt disease-free soils (37). Hamid et al. demonstrated that *Mortierella* and *Purpureocillium* are related to soil suppressiveness against soybean cyst nematode (41). Additionally, the genus *Purpureocillium* is recognized as a promising biological control agent, effectively parasitizing the eggs of plant-parasitic nematodes (42). Lopez et al. found that the endophytic fungus *Purpureocillium lilacinum* enhances the growth of cultivated cotton (*Gossypium hirsutum*) and inhibits cotton bollworm (*Helicoverpa zea*) (43). Certain species within the genus *Gibellulopsis* are classified as pathogenic fungi, such as *Gibellulopsis nigrescens*, which causes wilt in sugar beets (44). However, numerous hypovirulent strains exist, including *Gibellulopsis nigrescens* Vn-1 and *Gibellulopsis nigrescens* CVn-WHg, which are employed as biological control agents against Verticillium wilt in sunflowers and cotton (45–46). Jin et al. also discovered that crop rotation increased the relative abundances of *Gibellulopsis* in the cucumber rhizosphere, indicating the beneficial role of the genus *Gibellulopsis* in healthy soils (47). The genus *Xenomyrothecium* is classified within the group of saprophytic fungi, which play a crucial role in the decomposition of plant biomass, highlighting their ecological significance (48). Furthermore, Sundar and Arunachalam discovered that *Xenomyrothecium tongaense* PTS8, a rare endophyte of *Polianthes tuberosa*, exhibits significant antagonism against multidrug-resistant pathogens (49).

Therefore, we postulated that strain SCA4-21 may synergistically control banana *Fusarium* wilt by producing antifungal compounds and enriching beneficial bacteria and fungi. The functions of bacteria and fungi still needs to be further explored in the decrease of *Fusarium* abundance in soil.

## 5. Conclusion

The extracts of *S. luomodiensis* SCA4-21 significantly inhibited hyphae growth and spore germination of Foc TR4, severely destroyed the ultrastructure of Foc TR4 hyphal cells and spore morphology, and exhibited excellent broad-spectrum antifungal activity against eight other phytopathogenic fungi. GC-MS analysis revealed that strain SCA4-21 produces a total of 32 volatile organic compounds, including the anticancer compound Butanoic acid methyl ester, the nematicidal compound Methyl isovalerate, the growth-promoting compound 3-Octanone, and the antifungal compounds 5-Methyl-2-heptanone, 6-Methyl-2-heptanone, Hexanoic acid methyl ester, and Benzoic acid methyl ester. Pot experiment revealed that the inoculation of strain SCA4-21 significantly inhibited infection of Foc TR4 corms of banana seedlings, resulting in a notable decrease in the disease index and an improvement in the growth of banana seedlings. High-throughput sequencing indicated that the inoculation of strain SCA4-21 significantly enhanced the population of beneficial microorganisms, including bacterial genera like *Streptomyces*, *Bacillus*, *Sphingomonas*, and *Massilia*, and fungal genera such as *Mortierella*, *Purpureocillium*, *Gibellulopsis*, and *Xenomyrothecium*, while significantly reducing pathogens such as the bacterial genus *Pantoea* and the fungal genus *Fusarium* in banana rhizosphere soil. Spearman bacterial correlation network analysis revealed that the genus *Streptomyces* exhibited a positive correlation with the genera *Bacillus*, *Sphingomonas*, *Massilia*, *Burkholderia*-*Caballeroni*a-*Paraburkholderia*, and *Cupriavidus*, while a negative correlation with the genus *Pantoea*. Tax4Fun2 prediction indicated that T3 group exhibited significant enrichment in pathways related to carbohydrate metabolism, amino acid metabolism, lipid metabolism, metabolism of terpenoids and polyketides, and biosynthesis of other secondary metabolites, suggesting that strain SCA4-21 could play an important role in carbon and nitrogen metabolisms within the ecological environment.

Therefore, *S. luomodiensis* SCA4-21 represents a potential bioagent for the control of banana *Fusarium* wilt. This bioagent may achieve this by producing antifungal compounds, enriching beneficial bacteria and fungi, and simultaneously reducing the abundance of *Fusarium*.

## Materials and methods

### 2.1 Extraction and purification of strain SCA4-21 extracts

The extracts of strain SCA4-21 were extracted and purified following the method of Li et al. (10). Namely, strain SCA4-21 was inoculated into 1000 mL of SLM (20 g of soluble starch, 15 g of soybean powder, 5 g of yeast powder, 2 g of peptone, 4 g of CaCO_3_, 4 g of NaCl, 1 L of ddH2O, pH 7.2–7.4) and cultured at 28°C for 7 days at 180 r / min. Then the fermentation broth was added 95% ethanol in a 1:1 ratio and shocked at 28 °C and 180 r / min for 2 days. The mixtures were filtered by filter paper, and the filtrate was concentrated by rotary vacuum evaporator (N-1300, BoLang Co., Ltd., Shanghai, China). To separate and purify extracts, the concentrate was fully adsorbed onto macroporous resin (D101) and eluted sequentially with 50%, 60%, 70%, 80% and 100% methanol gradient. These eluents were evaporated dry using rotary vacuum evaporator, respectively. These separated and purified extracts were dissolved in 10% of dimethyl sulfoxide (DMSO) with 20 mg/mL of a concentration, filtered using a 0.22-μm sterile filter (Millipore, Bedford, MA, United States) and preserved at -4 ℃, respectively.

The antifungal activity of the extracts against Foc TR4 was determined using agar dilution method (50). Briefly, 100 μL of extract solutions from different eluents were added to 60 mL of PDA medium that had been sterilized but not yet solidified, respectively. Every 60mL of PDA medium is evenly poured into three petri dishes to make plates. Three plates with 10% of DMSO were used as controls. Each plate in the center was inoculated a Foc TR4 mycelial plug with 0.5 cm diameter and incubated at 28 ℃ for 7 days. The clonal diameter of Foc TR4 was measured and fungal growth inhibition (FGI) was calculated using the following formula: FGI = [(C-T)/C] ×100%, where C and T represented the growth diameters of the control and the treatment group, respectively. The most effective extracts were selected for further investigations.

### 2.2 Measuring the EC_50_ of strain SCA4-21 extracts

PDA plates with final concentration extracts of 0.78, 1.56, 3.13, 6.25, 12.50, 25.00, 50.00, 100.00, 200.00 μg/ml respectively were made up. The fungal disk of Foc TR4 (0.5cm diameter) was inoculated on PDA plates and cultivated in an incubator at 28℃ for 7 days. The plates with DMSO (10%, v/v) were used as a control. Three biological replicates for each concentration were performed. The growth diameter of Foc TR4 was measured, and the percentage of mycelial inhibition was calculated. A linear regression equation was established using the least square method to determine the half maximal effective concentration (EC50) of extracts on the mycelial growth of Foc TR4 (16).

### 2.3. Determining broad-spectrum antifungal activity of strain SCA4-21 extracts

The broad-spectrum anttifungal activity of strain SCA4-21 extracts against eight plant pathogenic fungi were determined using agar well diffusion method (51). The eight pathogenic fungi include *Curvularia fallax* (ATCC 34598), *Colletotrichum gloeosporioides* (Penzig) Penzig et Saccardo (ATCC MYA-456), *Colletotrichum gloeosporioides* (ACCC 36351), *Colletotrichum acutatum*(ATCC 56815), *Colletotrichum fragariae* (ATCC 58718), *Colletotrichum gloeosporioides (Penz) Saec* (ACCC 36351), *Fusarium oxysporum* f. sp. *cucumerinum* Owen. anamorph (ATCC 204378), *Fusarium graminearum* (DSM 21803). The pathogens were provided friendly by the Institute of Environment and Plant Protection, China Academy of Tropical Agricultural Sciences, Haikou, China. Fungal growth inhibition (FGI) was calculated as above formula.

### 2.4. Effect of strain SCA4-21 extracts on spore germination of Foc TR4

Effect of strain SCA4-21 extracts on spore germination of Foc TR4 was observed using our previous method (52) with minor modifications. 5000 mL of sterile water was added to the well-growing Foc TR4 plate, and the mycelia and spores were scraped using the sterilized coating rod. The spore suspension obtained by filtering off mycelia was diluted to 1×10^6^ CFU / ml and mixed with equal volume of different concentration extracts (1 × EC_50_, 2 × EC_50_, 4 × EC_50_, 8 × EC_50_). 10% DMSO was used as control instead of extracts. After incubate at 28℃ for 24h, The spore germination of Foc TR4 was observed by an inverted microscope (Cellcutplus, MMI, Germany), 100 conidia of each field were counted. The spore germination inhibition (SGI) was calculated following the formula: SGI = (A -B)/A×100%, where A and B signify the spore germination rate of the control group and the treatment group, respectively.

#### Effect of strain SCA4-21 extracts on mycelial and spore morphology as well as the ultrastructure of Foc TR4

Effect of strain SCA4-21 extracts on mycelial morphology of Foc TR4 was observed following the method of Cao et al. (53), with some modification. PDA plates containing 4×EC_50_ extracts of strain SCA4-21 were utilized as treatments, while PDA plates containing 10% DMSO served as controls. Disks (0.5cm-diameter) of Foc TR4 were inoculated at the centers of these plates. After 3 days of culture, fungal mycelia blocks (0.5-cm diameter) were cut from the edge of pathogen clone, fixed with 2.5% (v/v) glutaraldehyde (C_3_H_8_O_2_) at -4 ℃ overnight, and rinsed twice using phosphate-buffered saline (0.1 mol/L, pH 7.4). Then, those fungal mycelia blocks underwent gradient dehydration in 30%, 50%, 70%, 80%, 90%, 95% and 100% ethanol for 2 min each, followed by a rinse with tert-butanol for 20 min. These fungal mycelium blocks were subsequently soaked in fresh tert-butanol and frozen at -80 ℃. They were then subjected to freeze-drying using a freeze dryer. (FDU-2110, EYELA, Tokyo, Japan). The sample sheet was fixed on a small steel column and coated with a film of gold-palladium alloy under vacuum. Then the morphology of the hypha was observed by scanning electron microscopy (SEM, Zeiss ∑IGMA, Germany). Fungal hyphae were also collected from the above 3-day-old treatment and control plates, respectively and treated with according to our previous method (53). The ultrastructure of fungal hyphae was detected by transmission electron microscopy (JEM-1400Flash, Japan).

In addition, Foc TR4 spores suspension with 1×10^6^ CFU concentration was prepared according to above method and mixed with 4×EC_50_ extracts solution of strain SCA4-21 (v:v=1:1). 20 μL of mixture was dropped onto a cover plate which was placed on a glass slide and cultivated for at 28 ℃ 24 h. Control was set up by replacing extracts solution with 10% DMSO. Spores were treated as described above mothed and observed by SEM (Zeiss ∑IGMA, Germany).

#### Identification of the volatile organic compounds of strain SCA4-21

To identify the volatile organic compounds of strain SCA4-21, one hundred microliters of strain SCA4-21 suspension (10^6^ CFU mL^-1^) was inoculated into a sterilized 50-mL flask containing 15 mL of SLM. Equal volume of ddH_2_O replacing strain SCA4-21 suspension was used as control. Both treatment and control were set up with three replicates each. After being cultured for 7d at 28℃ with agitation at 180 rpm, the volatile organic compounds of strain SCA4-21 were analyzed using GC-MS (Clarus690+SQ8T, PerkinE1mer, USA) following the method described in our previous study (53) and identified by comparing the mass spectra of treatments with those of controls.

#### Evaluation of the biocontrol efficiency of strain SCA4-21 and its promotion effect on banana seedlings

The pot experiment of banana seedlings was carried out to assess biocontrol potential of strain SCA4-21 against banana fusarium wilt. GFP-Foc TR4 strain expressing green fluorescent protein (GFP) was provided kindly by the Institute of Tropical Bioscience and Biotechnology, China Academy of Tropical Agricultural Sciences, Haikou, China. The spore suspension of GFP-Foc TR4 was produced by inoculating the pathogen in potato dextrose broth medium (PDB) and shaking (180 rpm) at 28◦C for 7 days, and adjusted to final concentration of 1×10^6^ CFU/mL. The roots of healthy banana seedings with 3-4 leaves were wound with scissors and immersed in the spore suspension of GFP-Foc TR4 for 30 min, then the seedings were planted in plastic pots filled with 900 g of soils to use as positive control or treatment. Healthy banana seedings untreated with GFP-Foc TR4 were planted as negative control. Namely, three experimental groups were designed: T1 (sterilized soybean liquid medium: SLM), T2 (Foc TR4+ SLM) and T3 (Foc TR4+SCA4-21). Each treatment group is 30 pots. The fermentation broth of strain SCA4-21 derived from inoculating the isolates into SLM and shaking (180 rpm) at 28◦C for 7 days. T1 group seedlings were watered with 100 mL of SLM diluted 10 times once a week. T2 and T3 group seedlings were irrigated with spore suspension of GFP-Foc TR4 at concentration of 1×10^6^ CFU/mL and 100 ml of SLM or fermentation broth diluted 10 times once per week.

After banana seedlings planted in a greenhouse at 28℃, with 70% of humidity and 12 h dark/12 h natural light for 60 days, the corms and roots of the banana seedlings were cut to observe the infection degree of Foc TR4 using laser confocal microscope (Axio Scope A1, Carl ZEISS, Germany). The physiological and biochemical indexes of banana plants under experimental group were determined, including chlorophyll content, leaf area, plant height, stem diameter, and biomass (dry weight, fresh weight) at 120 th day. The disease index and biocontrol efficiency were calculated according to the method of Chen et al. (15).

#### Effect of strain SCA4-21 on the microbial community structure of banana rhizosphere soil

To decipher the effect of strain SCA4-21 on the microbial community structure of banana rhizosphere soil, a total of 3 rhizosphere soil samples were collected for each experimental group and stored in a -80℃. The collected samples were sent to Beijing Baimaike Biotechnology Co., Ltd. for microbial diversity sequencing using paired- end method on Illumina NovaSeq sequencing platform. Bioinformatics analysis was performed using BMKCloud (www.biocloud.net). Raw reads were filtered using the trimmomatic (v 0.33) software (54) and adapter sequences were removed to acquire clean reads using cutadapt (v1.9.1) (55). Clean reads were denoised, spliced and removed the chimeric sequences using the dada2 method in QIIME2 (v 2020.6) software to obtain non-chimeric reads (56–57). Operational taxonomic units (OTUs) were identified and Venn diagrams were made using QIIME2 (v 2020.6) software (57). The bacterial and fungal alpha and beta diversity indices were analyzed using QIIME2 (v 2020.6) (57). Specifically, Shannon, Simpson, Chao1 and ACE indices were measured to estimate microbial alpha diversity. The Wilcoxon test was selected for between-group variance analysis of alpha diversity indices at OUT level. Principal components analysis (PCA) and nonmetric multidimensional scaling (NMDS) were visualized to assess microbial beta diversity. The Binary-Jaccard distance metric was chosen for analyzing between-group variance in the NMDS analysis. Taxonomic annotation of OTUs was performed using SILVA database (58) (http://www.arb-silva.de/). A community structure map at the level of microbial genera was created using R language tools in QIIME2 (v 2020.6) software (57). Intergroup differences in abundance of microbial genera levels were analyzed using analysis of variance (ANOVA) method. Functional profiles of bacteria community were predicted using Tax4Fun2 (57). Bacterial network analysis was performed using Molecular Ecological Network Analyses Pipeline to elucidate the effects of different treatments on bacterial ecological network (60) (http://ieg4.rccc.ou.edu/mena).

#### Statistical Analysis

Statistical analysis was performed with the SPSS Version 22.0 software (SPSS Inc., Chicago, IL, United States). Significant difference between means was determined by the Duncan’s multiple range test at p < 0.05. The difference among treatments was determined using one-way analysis of variance (ANOVA). All data from three biological replicates of each experiment were obtained as means ± the standard error (SE).

## Data Availability

The data that has been used is confidential.

## Acknowledgments

This study was supported by the project of National Key Laboratory for Tropical Crop Breeding (NKLTCB202306), the National Natural Science Foundation of China (U22A20487 and 322MS126), The Natural Science Foundation of Hainan 322QN417 and 322MS126), Chinese Academy of Tropical Agricultural Sciences for Science and Technology Innovation Team of National Tropical Agricultural Science Center (CATASCXTD202309 and CATASCXTD202312), Central Public-interest Scientific Institution Basal Research Fund (1630052023011) and the China Agriculture Research System (CARS-31).

---

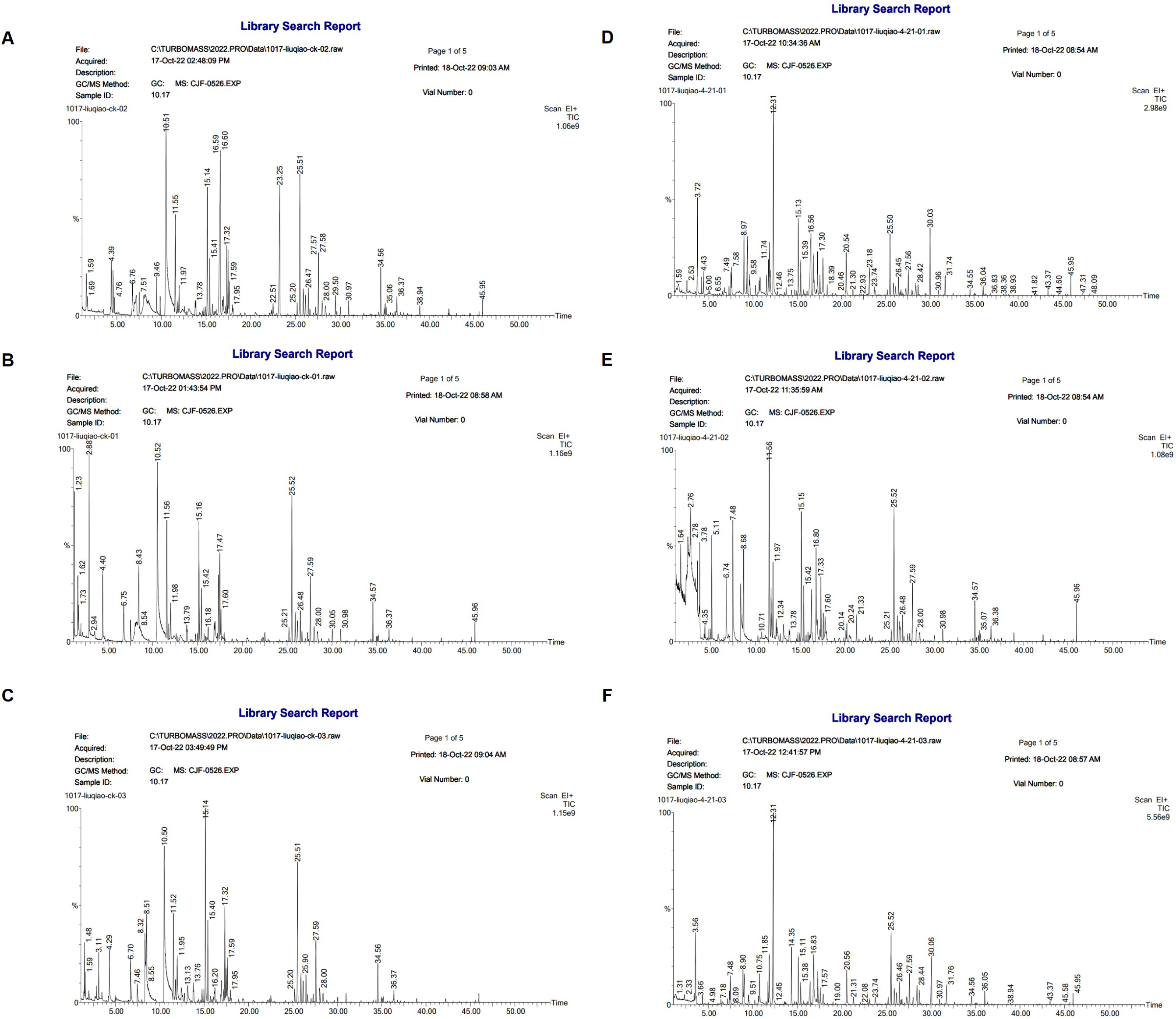

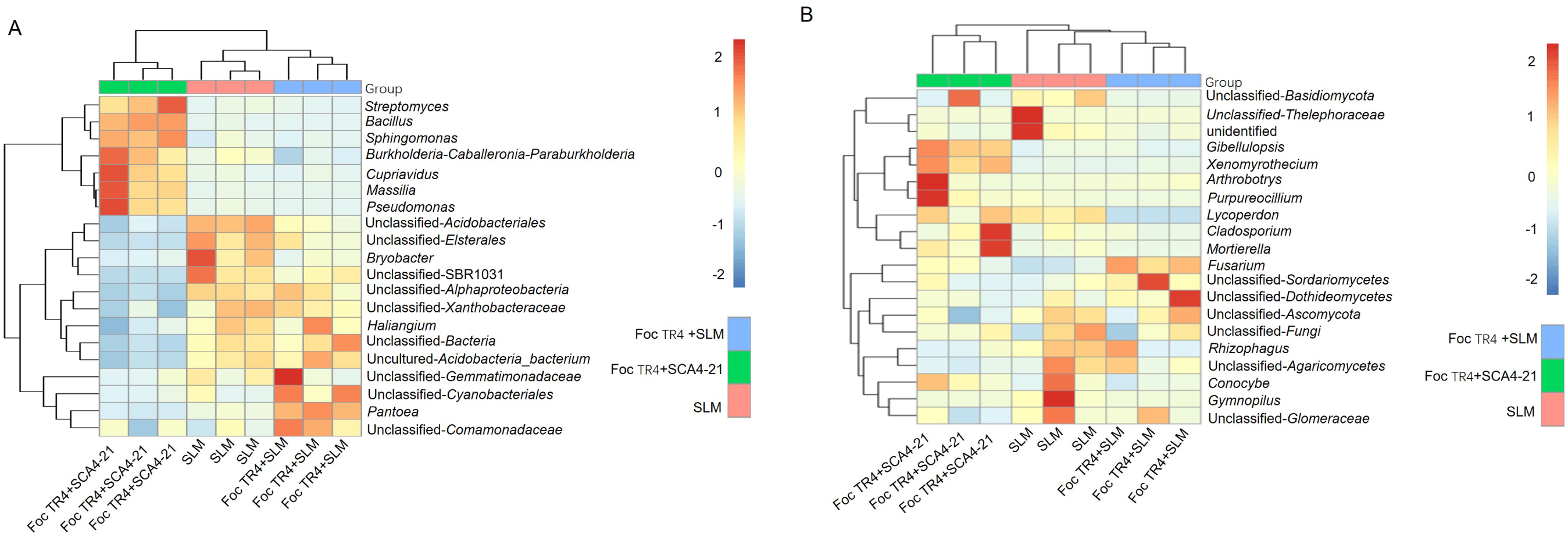

